# Comparisons of Coding and Noncoding Sequences to infer the Origin of Codon usage Bias

**DOI:** 10.1101/174359

**Authors:** Prashant Mainali, Sobita Pathak

## Abstract

Codon usage bias is the preferential use of the subset of synonymous codons during translation. In this paper, the comparisons of normalized entropy and GC content between the sequence of coding regions of *Escherichia coli k12* and noncoding regions (ncRNA, rRNA) of various organisms were done to shed light on the origin of the codon usage bias.The normalized entropy of the coding regions was found significantly higher than the noncoding regions, suggesting the role of the translation process in shaping codon usage bias. Further, when the position specific GC content of both coding and noncoding regions was analyzed, the GC2 content in coding regions was lower than GC1 and GC2 while in noncoding regions, the GC1, GC2, GC3 contents were approximately equal. This discrepancy is explained by the biased mutation coupled with the presence and absence of selection pressure. The accumulation of CG content occurs in the sequences due to mutation bias in DNA repair and recombination process. In noncoding regions, the mutation is harmful and thus, selected against while due to the degeneracy of codons in coding regions, a mutation in GC3 is neutral and hence, not selected. Thus, the accumulation of GC content occurs in coding regions, and thus codon usage bias occurs.

## INTRODUCTION

The genetic code decides which amino acid will be inserted during translation of protein for each three nucleotide set, or codon, in the mRNA. [1] More precisely, the genetic codon determines which of the 61 triplets, or codon, corresponds to which of the 20 amino acids. [2] There are 64 possible codons – three codons terminate translation while remaining 61 codes for amino acids, but only 20 different amino acids to incorporate during translation. Thus, codons are redundant, and more than one codon encodes the same amino acids. This feature of the genetic code is mentioned as the degeneracy of the codon. [3] Although few amino acids are encoded by a single codon, most amino acids are encoded by two to six different codons. Different codons that encode the same amino acid are known as synonymous codons. [2]

The frequency of synonymous codons is not fair as the determination of a large number of DNA sequences from different species has shown, in many cases, that synonymous codons for any amino acids are not used randomly. [4] This suggests that the sequence in the DNA is not random, but some bias occurs. Some synonymous codons are highly expressed, whereas the use of others is limited. This phenomenon of preferential use of the subset of synonymous codons during translation is known as codon usage bias.

If during a translation of a gene, a synonymous codon incorporates a particular amino acid more often, a gene is said to have codon usage bias. Such frequently used codons are known as optimized codons.

The codon usage bias differs among organisms. Closely related organisms have similar patterns of codon usage. [5] The genome hypothesis of codon usage bias states that different organisms have distinct codon biases [6]. Even the difference in codon usage is observed among genes of the same organism. It has been revealed through multivariate analyses that even in each species codon usage among genes differ. [5] The codon usage of sixteen weakly and thirteen strongly expressed *E. coli* genes were examined, yet again using a multivariate analysis technique, and was found to have a noticeable variation in codon usage [7]

It was proposed that codon usage bias might be correlated with variation in tRNA abundance [7].The codon usage bias in *E. coli* and *Saccharomyces cerevisiae* was correlated with the abundance of the cognate tRNA. [8, 9]The optimal codons are one with the highest cognate tRNA. The experiments performed by Sorensen *et al* (1989)have detected differences in these rates.[10] Codons that are recognized by the major tRNAs are translated 3–6 fold faster than their synonyms This selection may favor the use of the more frequent codons during translation.[10] Additionally, the experiments performed by Parjer et al. (1983) has estimated that in *E. coli* the non-optimal Asparagine codon (AAU) can be mistranslated eight to ten times more often than its optimal synonym (AAC). [11] These findings imply that the codon usage bias or use of optimal codon for translation increases the accuracy of translation with fewer mistakes than genes that use less frequent codons and also increases the speed of translation. Hence, the organism may confer selective advantage due to the use of optimal codons, and codon usage bias is maintained. Therefore, translational selection can be a responsible factor for shaping codon usage bias.

Moreover, highly expressed genes generally show a higher bias in synonymous codon usage. The positive correlation between codon usage bias and expression levels is again attributed to selection for translational efficiency. [12] It is also showed that codon usage bias modulates gene expression. [7]

Although translational selection plays a role in shaping codon usage bias within species, such selection may not be directly correlated or explained with the codon usage bias pattern observed among species because much of the variation among species appears to be due to differing patterns of mutation. [13] An alternative explanation for the cause of codon usage bias, a mutational explanation, suggests that codon usage bias arises from the properties of underlying mutational processes – e.g. biases in nucleotides produced by point mutations, contextual biases in the point mutation rates, or biases in repair. [14] Mutational explanations are neutral, because they confer no fitness advantage or detriment associated with alternative synonymous codons. Mutational mechanisms are typically invoked to explain inter-specific variation in codon usage, especially among unicellular organisms. [15]

Recombination also affects codon usage bias. The study performed by Comeron et al. (1999) observed a positive correlation between recombination rate and codon usage bias. [16] Other factors that may influence codon usage bias include: GC content [16, 17]; biased gene conversion. [18, 19]

It is evident from above studies that the codon usage bias is commonly observed the phenomenon in the genome, be it inter-genome or intra-genome, but the origin of codon usage bias is a complex puzzle that is rarely explained by only one factor.

The different quantitative methods are available to quantify or characterize codon usage bias. The codon preference bias, codon bias index (CBI), scaled Chi-square approach, codon preference statistic (CPS), relative synonymous codon usage (RSCU), codon adaptation index (CAI), effective number of codons (ENC), Shannon information theory, synonymous codon usage order (SCUO) are the available quantitative methods to characterize codon usage bias in terms of numbers. [20] Each method has own strength and limitations.

After Shannon published a pioneering paper on “A Mathematical Theory of Communication” [21], the use of information theory is not limited to telecommunication and data compression but has been used in many fields of physics, biology, and linguistics. In recent times, the use of Shannon’s entropy in molecular biology for prediction of DNA, RNA structure, protein secondary structure prediction, modeling molecular interaction, drug design has been recommended. [22] In this paper we have used Shannon informatics theory to measure codon usage bias in terms of Synonymous codon usage order (SCUO). The fundamental principle involved in this method is the calculation of the entropy of each synonymous codon for each amino acid in the protein.

## MATERIALS AND METHODS

### Materials

The 20 DNA sequences of coding regions of Escherria coli k12 and 20 DNA sequences of non-coding regions (ncRNA, rRNA) of various organisms were obtained in FASTA format in NCBI database.

### Methods

### Calculation of SCUO

The information theoretic value of a given DNA sequence was obtained using the Shannon formula [23].

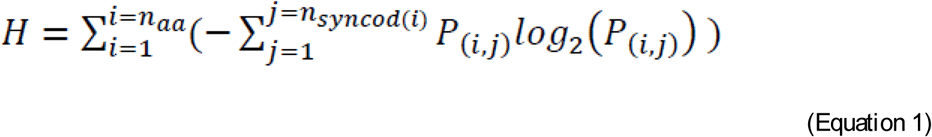

Here,

n_aa_ is the number of distinct amino acids,
n_syncod(i)_ is the number of synonymous codons for each amino acid *i* (or macro-state) whose value range from 1 to 6,
P_(i,j)_ is the probability of synonymous codon *j* for amino acid *i*.

For every sequence, the probability of each synonymous codon was calculated using online sequence manipulation suite using the standard genetic code as the reference. [24] The probability of synonymous codons was feed into (Equation 1) to calculate the entropy. This was achieved using the tailor made program in Python. The obtained value of H is the entropy per codon of the gene.

Additionally, to calculate the maximum entropy of the sequence, a random sequence of the same length as that of a gene to be compared and have the likelihood of nucleotides 25%, was created using online platform. [25] Similarly, the probability of each synonymous codon was calculated and hence, maximum entropy per codon of the random sequence is calculated using the python program.

Now, subtracting observed entropy with the maximum entropy gives information which measures the non-randomness in synonymous codon usage and hence describes the degree of orderliness for synonymous codon usage in each sequence.

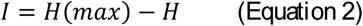

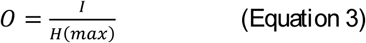

O is the normalized difference between the maximum entropy and the observed entropy in the sequence. 0 ≤ O ≤ 1. When O=1, usage is biased to the extreme while O=0, usage is random. The O is designated as the synonymous codon usage order (SCUO). [26]

The normalized difference between the maximum entropy and the observed entropy in the sequence i.e. normalized entropy was calculated, for both the noncoding and coding sequences and was compared.

### Calculation of position specific GC content

The position specific GC content of noncoding and coding sequence was calculated using the python program. The GC content of noncoding and coding sequences was compared.

## RESULTS AND DISCUSSION

The normalized entropy of coding regions of *Escherichia coli k12* and 20 DNA sequences of non-coding regions (ncRNA, rRNA) of various organisms was compared using independent t-test. The P value was less than significant level of 0.01; thus, the null hypothesis that says both mean are same was rejected and the alternate hypothesis that says the mean data are significantly different at 99 % confidence interval and a significant level of 0.01 was accepted. (Figure I)

**I. Figure:**
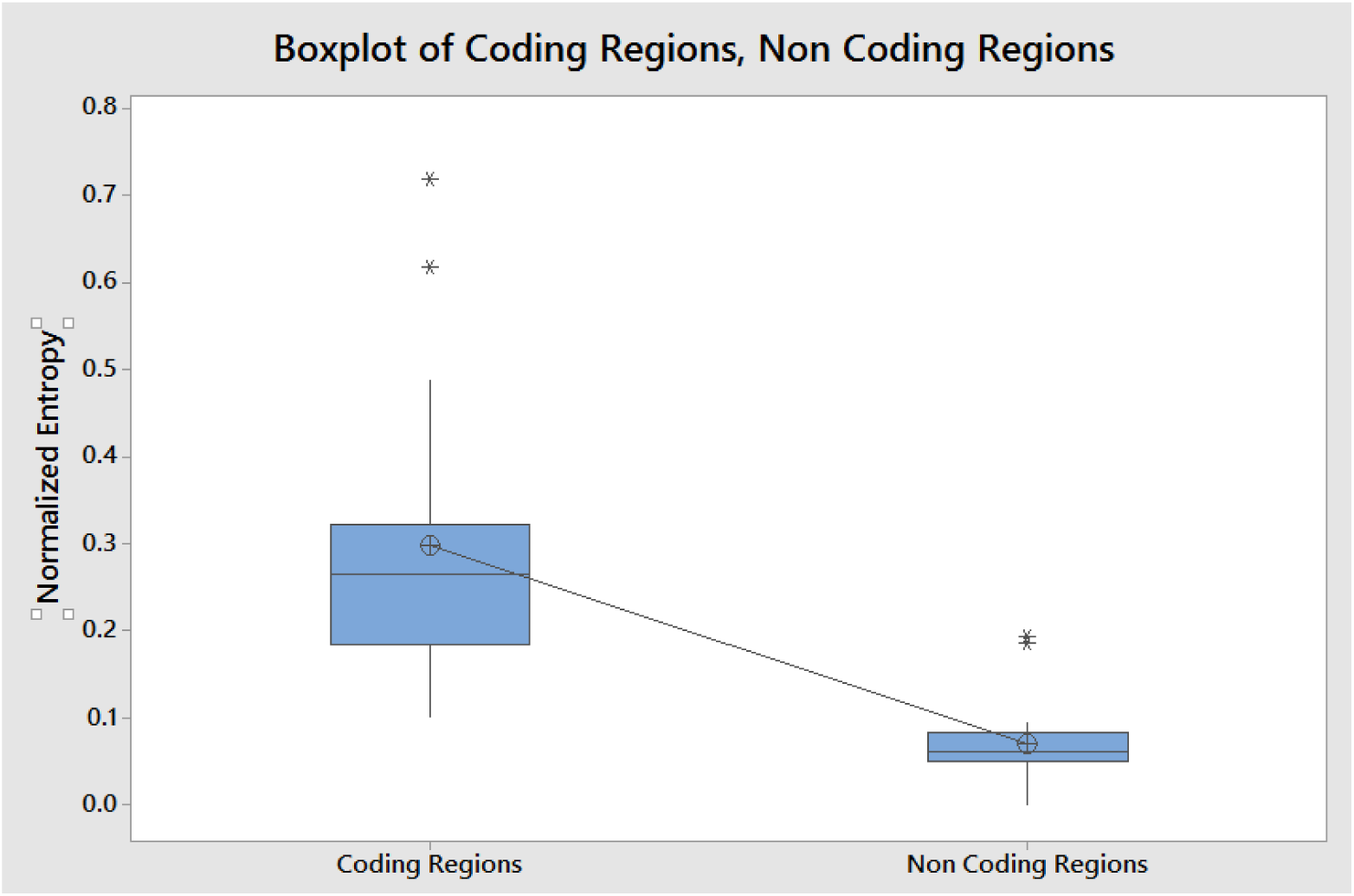
The result of independent t-test of normalized entropy of coding regions and noncoding regions.

Hence, from the results, it can be concluded, the normalized entropy of noncoding region was greater than the normalized entropy of coding regions. As the coding and non-coding regions were independently selected, we assume this result can be extrapolated for all the coding and non-coding regions. This infers, although, noncoding regions also show slight codon usage bias; the codon usage bias in coding regions is prominent. As, the non-coding regions do not go through the translational process while coding region goes through this, this difference in codon usage bias can be attributed to the translational process or due to some idiosyncrasy of the translational mechanism.

The role of the translation process in codon usage bias is established from above observation. The possibility arises that the translation can select the optimal codons. We know codons that are recognized by the major tRNAs are translated 3–6 fold faster than their synonyms. This may favor the use of the more frequent codons during translation. [10] However, it cannot be concluded that the translation selects optimal codons. We do not know the selection pressure generated by translation.

Further, the GC content in the genome is correlated with the codon usage bias. [16, 17] Thus, in this research, the normalized entropy and the GC content of the respective DNA sequences (both coding and noncoding) were analyzed and we found the weak correlation with the Pearson’s coefficient of 0.269. Due to the low amount of data, the correlation is probably an underestimate. But, the interesting thing is that the correlation between the normalized entropy and the position specific GC content i.e. GC1 and GC3 is higher than the correlation with the total GC content of the sequences. The Pearson’s correlation coefficient between the normalized entropy and the GC1 content is 0.496. Similarly, the Pearson’s correlation coefficient between the normalized entropy and the GC3 content is 0.359. The reason for higher correlation with GC1 content is not explained here. But, the higher correlation for GC3 content can explain the origin of codon usage bias.

Additionally, the next interesting fact was also deciphered from the comparison of the coding and noncoding sequences. In noncoding sequences, the position specific GC content (GC1, GC2, GC3) was approximately equal. (Figure II). Conversely, in noncoding sequences, the GC content varied. In all the cases, the GC1 and GC3 content were greater than GC2 content.(Figure III) The question arises why the noncoding sequences have equal position specific GC content while coding sequences do not hail that feature. There are biases for C and G nucleotides in coding sequences. The major driving factor for these biases can be explained by mutational bias. [27]The discrepancy of higher CG content in coding sequence can be explained by mutational bias and neutral theory of molecular evolution.

**II. Figure:**
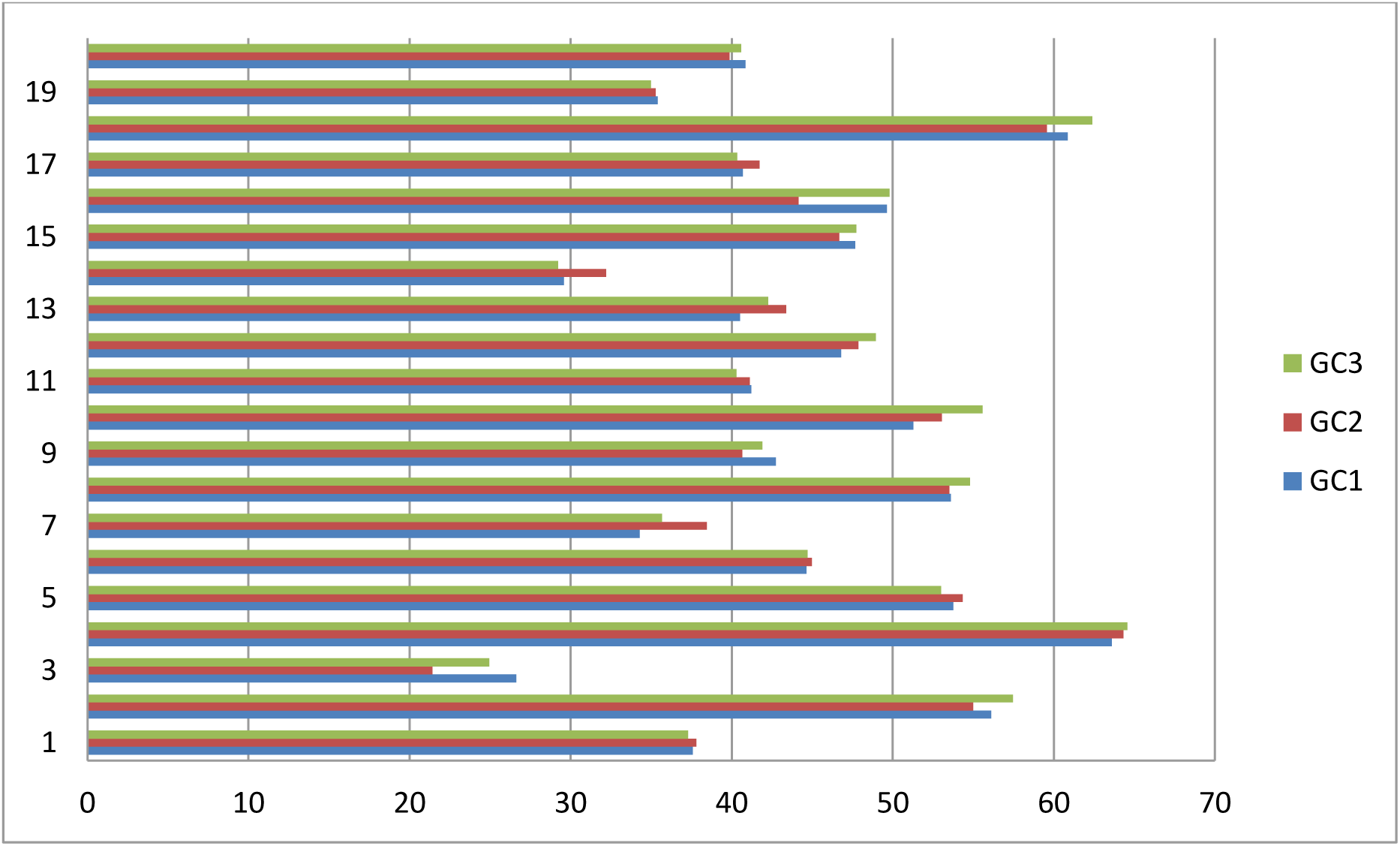
The position specific GC percentage of noncoding regions.

**III. Figure:**
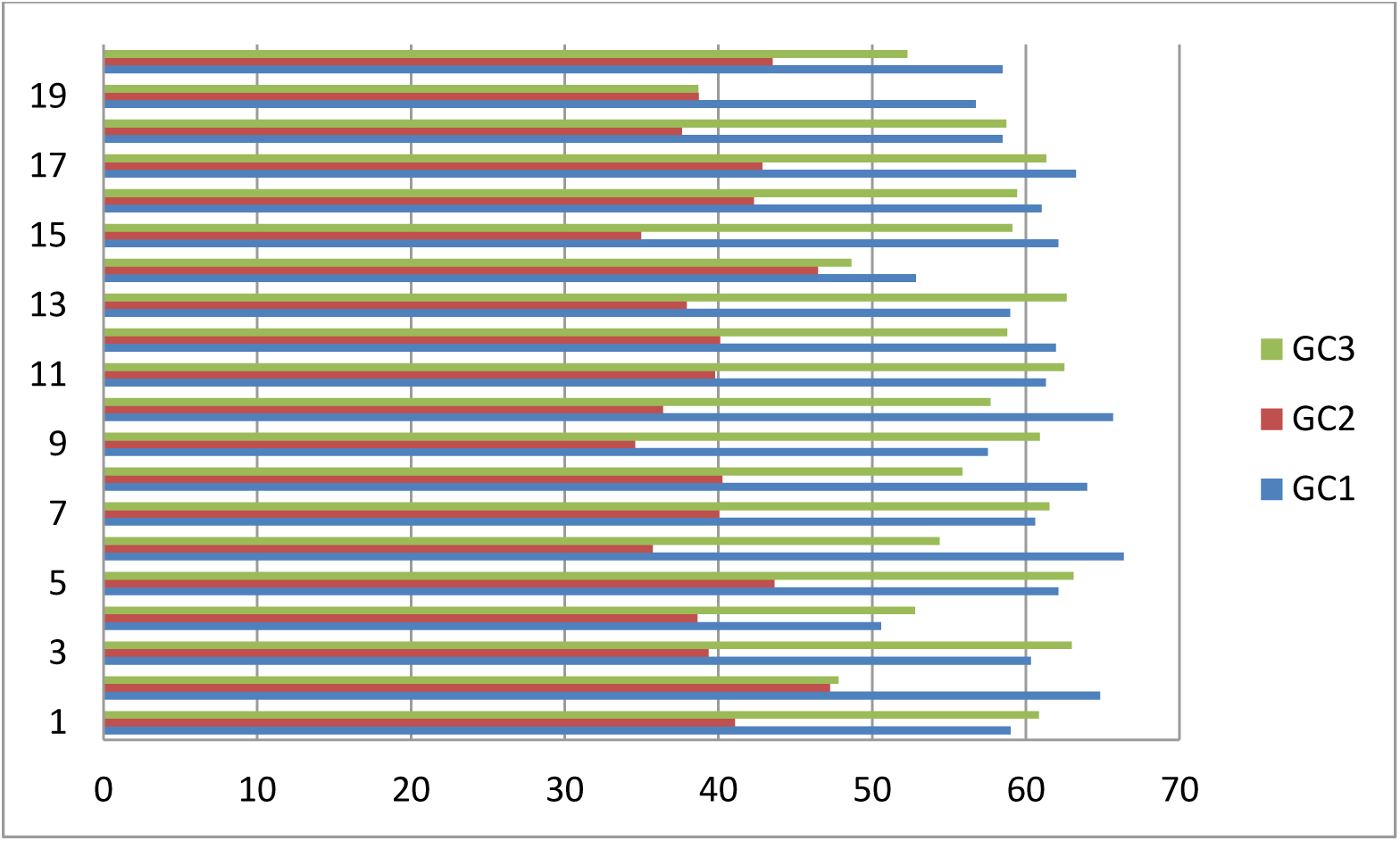
The position specific GC percentage of coding regions.

At equilibrium, the number of AT → GC and GC →AT mutations are equal. For biased mutations, the equilibrium is disturbed, and DNA sequence accumulates a pair of nucleotides in greater quantity. In coding sequences, the biased mutation was pronounced in GC3 (the reason why GC1 also has more GC content is not explained). There is experimental evidence that demonstrates that the DNA repair mechanism and recombination mechanism has biases towards AT→CG mutations. [27] Gerton *et al.* have found the correlation between the recombination and GC content in *Saccharomyces cerevisiae*[28] while Marais *et al.* 2001 has found the correlation between the recombination and GC content in *Drosophila melanogaster*, and *Caenorhabditis elegans.*[17] Similarly, Brown and Jiricny 1987; Bill *et al.* 1998 have experimentally observed biased DNA repair toward GC in mammalian cells after transfection of mismatched DNA fragments. [29, 30]So, it is likely that DNA sequence accumulates more CG. The high GC3 content in the coding regions only but not in noncoding sequence can be explained by the neutral theory of evolution. The neutral theory of molecular evolution, proposed by Kimura, states that the vast majority of conserved mutations are neutral. [31] Similarly, in the histrionic paper “Codon—anticodon pairing: The wobble hypothesis” Crick has suggested that due to the degeneracy of the genetic code, the first two positions of the codon or triplet are important for pairing while there may be some wobble in the pairing of the third base. [32]To simplify, the third base is considered to be less discriminatory than the other two bases. This implies that the third base of the codon is subjected to no or less selection pressure if a mutation happens there. The mutation in the third base of the codon does not change the amino acid profile of the protein. Thus, the mutation in the third base of the triplet is neutral. This helps in the accumulation of the high GC3 content in the coding sequences as selection against this mutation doesn’t occur. Similarly, the biases are not observed in the noncoding sequence because the mutation in the noncoding sequence is not neutral. The mutations in any position in the noncoding sequence (ncRNA, rRNA) alter the structure of this RNA, hence, altering the function of these RNA. [1]So, the selection occurs against mutations in noncoding sequences. The accumulation of the CG in the third base of coding sequence ensue high amount of codons ending in C&G and low number of codons ending with A&T, thus, demonstrating codon usage bias. The codon usage bias will ultimately benefit the cell as the optimal codons will translate the protein in higher rates. [10]

## CONCLUSION

The role of the translational role in shaping codon usage bias cannot be justified by this research. But, this research lights on the role of translational process on shaping codon bias. The DNA repair mechanism and recombination mechanism increases the CG content volume by the bias mutation toward CG. The translational foible is such; the mutation in the third base of sequence doesn’t alter the structure of the protein. Thus, if the bias mutation in GC3 occurs, the mutation is not selected but preserved. Hence, the GC3 content is higher in the coding regions. However, the selection is tough in the noncoding region (ncRNA, rRNA) and thus, mutations in the noncoding region is selected ensuing the equal distribution of GC content over the entire sequence. Hence, the biased mutation and the absence of selection pressure in the sequence are responsible for the codon usage bias in the coding regions.

## ACKNOWLEDGEMENT

We want to thank Mr. Aakash Pandey, for his work entitled ―Entropy and Codon Bias in HIV-1” inspired us to work in this research. Further, he provided us with the knowledge that provided us enough momentum to work in this research. We are indebted to Mr. Aryan Neupane for providing us with some valuable suggestions.

## TABLES

**I. Table:**
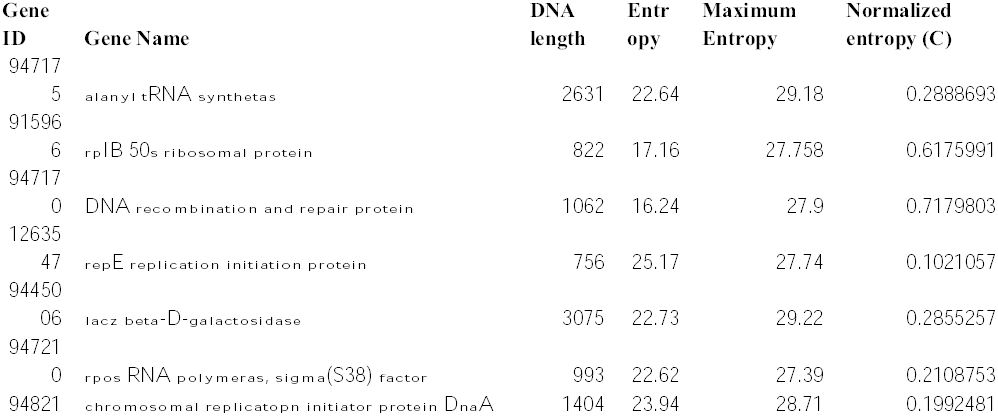
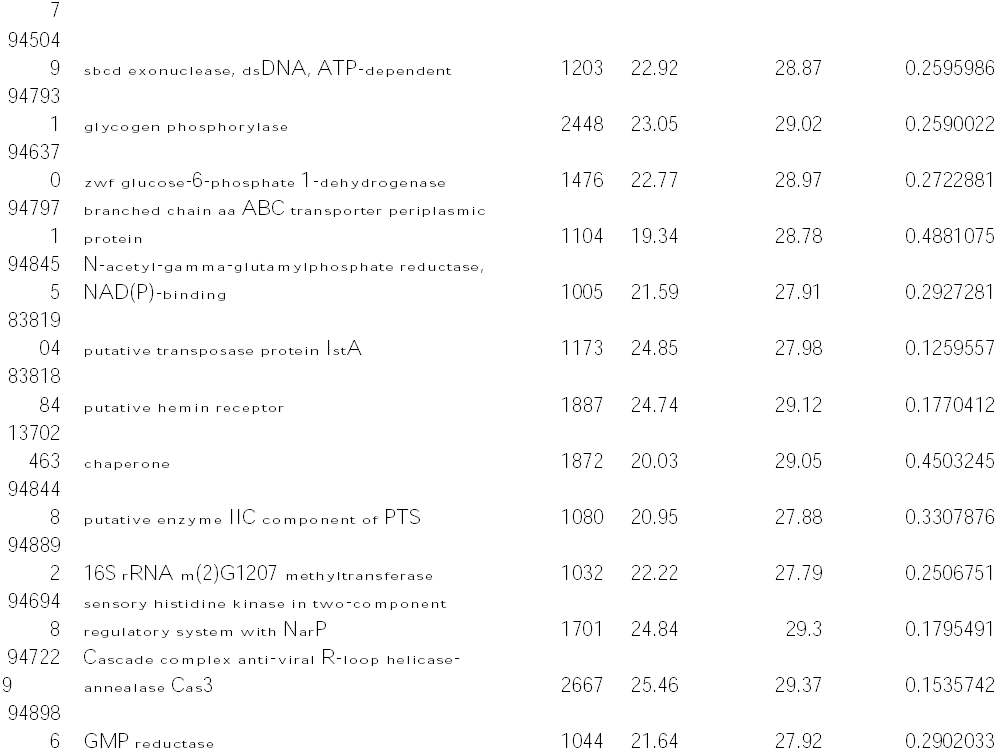
The normalized entropy of coding sequence

**II. Table:**
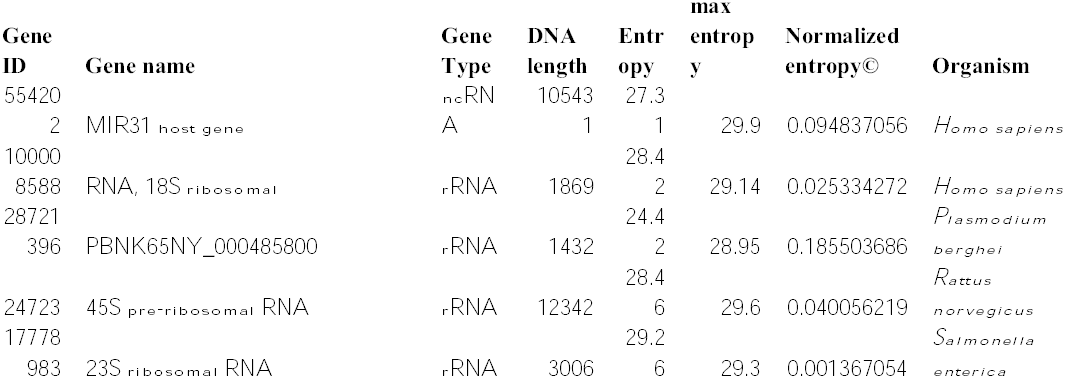
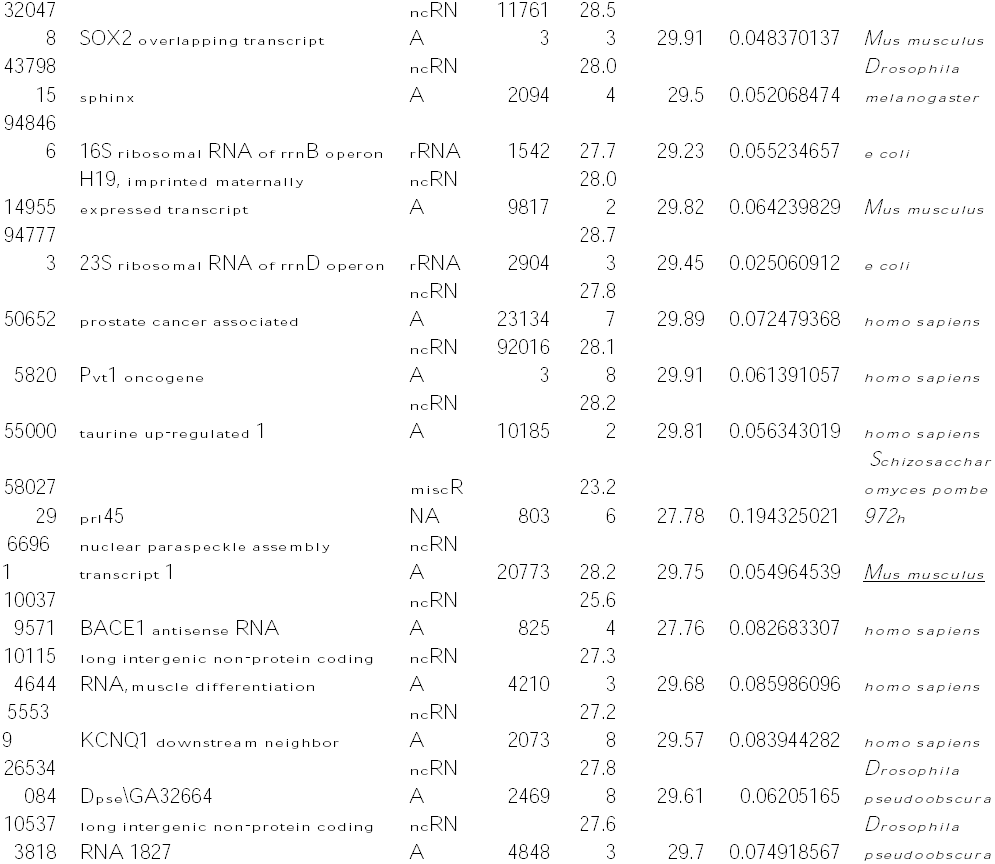
The normalized entropy of noncoding sequence

| Gene ID | Gene name | Gene Type | DNA length | Entropy | max entropy | Normalized entropy© | Organism |
| --- | --- | --- | --- | --- | --- | --- | --- |
| 55420 |  | ncRN | 10543 | 27.3 |  |  |  |
| 2 | MIR31 host gene | A | 1 | 1 | 29.9 | 0.094837056 | *Homo sapiens* |
| 10000 |  |  |  | 28.4 |  |  |  |
| 8588 | RNA, 18S ribosomal | rRNA | 1869 | 2 | 29.14 | 0.025334272 | *Homo sapiens* |
| 28721 |  |  |  | 24.4 |  |  | *Plasmodium* |
| 396 | PBNK65NY_000485800 | rRNA | 1432 | 2 | 28.95 | 0.185503686 | *berghei* |
|  |  |  |  | 28.4 |  |  | *Rattus* |
| 24723 | 45S pre-ribosomal RNA | rRNA | 12342 | 6 | 29.6 | 0.040056219 | *norvegicus* |
| 17778 |  |  |  | 29.2 |  |  | *Salmonella* |
| 983 | 23S ribosomal RNA | rRNA | 3006 | 6 | 29.3 | 0.001367054 | *enterica* |
| 32047 |  | ncRN | 11761 | 28.5 |  |  |  |
| 8 | SOX2 overlapping transcript | A | 3 | 3 | 29.91 | 0.048370137 | *Mus musculus* |
| 43798 |  | ncRN |  | 28.0 |  |  | *Drosophila* |
| 15 | sphinx | A | 2094 | 4 | 29.5 | 0.052068474 | *melanogaster* |
| 94846 |  |  |  |  |  |  |  |
| 6 | 16S ribosomal RNA of rrnB operon | rRNA | 1542 | 27.7 | 29.23 | 0.055234657 | *e coli* |
|  | H19, imprinted maternally | ncRN |  | 28.0 |  |  |  |
| 14955 | expressed transcript | A | 9817 | 2 | 29.82 | 0.064239829 | *Mus musculus* |
| 94777 |  |  |  | 28.7 |  |  |  |
| 3 | 23S ribosomal RNA of rrnD operon | rRNA | 2904 | 3 | 29.45 | 0.025060912 | *e coli* |
|  |  | ncRN |  | 27.8 |  |  |  |
| 50652 | prostate cancer associated | A | 23134 | 7 | 29.89 | 0.072479368 | *homo sapiens* |
|  |  | ncRN | 92016 | 28.1 |  |  |  |
| 5820 | Pvt1 oncogene | A | 3 | 8 | 29.91 | 0.061391057 | *homo sapiens* |
|  |  | ncRN |  | 28.2 |  |  |  |
| 55000 | taurine up-regulated 1 | A | 10185 | 2 | 29.81 | 0.056343019 | *homo sapiens* |
|  |  |  |  |  |  |  | *Schizosacchar* |
| 58027 |  | miscR |  | 23.2 |  |  | *omyces pombe* |
| 29 | prl45 | NA | 803 | 6 | 27.78 | 0.194325021 | *972h* |
| 6696 | nuclear paraspeckle assembly | ncRN |  |  |  |  |  |
| 1 | transcript 1 | A | 20773 | 28.2 | 29.75 | 0.054964539 | <u>*Mus musculus*</u> |
| 10037 |  | ncRN |  | 25.6 |  |  |  |
| 9571 | BACE1 antisense RNA | A | 825 | 4 | 27.76 | 0.082683307 | *homo sapiens* |
| 10115 | long intergenic non-protein coding | ncRN |  | 27.3 |  |  |  |
| 4644 | RNA,muscle differentiation | A | 4210 | 3 | 29.68 | 0.085986096 | *homo sapiens* |
| 5553 |  | ncRN |  | 27.2 |  |  |  |
| 9 | KCNQ1 downstream neighbor | A | 2073 | 8 | 29.57 | 0.083944282 | *homo sapiens* |
| 26534 |  | ncRN |  | 27.8 |  |  | *Drosophila* |
| 084 | Dpse\GA32664 | A | 2469 | 8 | 29.61 | 0.06205165 | *pseudoobscura* |
| 10537 | long intergenic non-protein coding | ncRN |  | 27.6 |  |  | *Drosophila* |
| 3818 | RNA 1827 | A | 4848 | 3 | 29.7 | 0.074918567 | *pseudoobscura* |

**III. Table:** The overall and position specific GC content of coding regions

II. Table: The normalized entropy of noncoding sequence
| Gene ID | Gene Name | GC(%) | GC1(%) | GC2(%) | GC3(%) |
| --- | --- | --- | --- | --- | --- |
| 947175 | alanyl tRNA synthetas | 53.3 | 59.01 | 41.09 | 60.84 |
| 915966 | rpIB 50s ribosomal protein | 53.1 | 64.83 | 47.25 | 47.81 |
| 947170 | DNA recombination and repair protein | 54.14 | 60.33 | 39.37 | 62.994 |
| 126354 | repE replication initiation protein | 47.22 | 50.59 | 38.64 | 52.78 |
| 7 |  |  |  |  |  |
| 944500 |  |  |  |  |  |
| 6 | lacZ beta-D-galactosidase | 56.26 | 62.1 | 43.65 | 63.12 |
| 947210 | rpos RNA polymeras, sigma(S38) factor | 52.06 | 66.36 | 35.75 | 54.38 |
| 948217 | chromosomal replicatopn initiator protein DnaA | 54.05 | 60.59 | 40.04 | 61.53 |
| 945049 | sbcd exonuclease, dsDNA, ATP-dependent | 53.28 | 64 | 40.25 | 55.86 |
| 947931 | glycogen phosphorylase | 50.9 | 57.54 | 34.6 | 60.9 |
| 946370 | zwf glucose-6-phosphate 1-dehydrogenase | 53.28 | 65.68 | 36.39 | 57.7 |
| 947971 | branched chain aa ABC transporter periplasmic protein | 54.6 | 61.3 | 39.78 | 62.5 |
|  | N-acetyl-gamma-glutamylphosphate reductase, NAD(P)-binding |  |  |  |  |
| 948455 |  | 53.53 | 61.97 | 40.11 | 58.8 |
| 838190 |  |  |  |  |  |
| 4 | putative transposase protein IstA | 53.11 | 58.97 | 37.94 | 62.65 |
| 838188 |  |  |  |  |  |
| 4 | putative hemin receptor | 49.28 | 52.86 | 46.49 | 48.64 |
| 137024 |  |  |  |  |  |
| 63 | chaperone | 52.02 | 62.11 | 34.99 | 59.13 |
| 948448 | putative enzyme IIC component of PTS | 54.16 | 61.02 | 42.33 | 59.44 |
| 948892 | 16S rRNA m(2)G1207 methyltransferase | 55.71 | 63.26 | 42.85 | 61.33 |
|  | sensory histidine kinase in two-component regulatory system |  |  |  |  |
| 946948 | with NarP | 51.61 | 58.48 | 37.63 | 58.73 |
| 947229 | Cascade complex anti-viral R-loop helicase-annealase Cas3 | 44.69 | 56.75 | 38.73 | 38.69 |
| 948986 | GMP reductase | 51.34 | 58.5 | 43.51 | 52.29 |

**IV Table:** The overall and position specific GC content of noncoding regions

III. Table: The overall and position specific GC content of coding regions
| Gene ID | Gene name | GC(%) | GC1(%) | GC2(%) | GC3(%) |
| --- | --- | --- | --- | --- | --- |
| 554202 | MIR31 host gene | 37.56 | 37.57 | 37.8 | 37.3 |
| 10000858 |  |  |  |  |  |
| 8 | RNA, 18S ribosomal | 56.12 | 56.1 | 54.98 | 57.46 |
| 28721396 | PBNK65NY_000485800 | 24.3 | 26.62 | 21.42 | 24.94 |
| 24723 | 45S pre-ribosomal RNA | 64.15 | 63.6 | 64.33 | 64.56 |
| 17778983 | 23S ribosomal RNA | 53.72 | 53.74 | 54.34 | 52.99 |
| 320478 | SOX2 overlapping transcript | 44.77 | 44.64 | 44.98 | 44.7 |
| 4379815 | sphinx | 36.1 | 34.28 | 38.45 | 35.67 |
| 948466 | 16S ribosomal RNA of rrnB operon | 54.4 | 53.6 | 53.5 | 54.8 |
| 14955 | H19, imprinted maternally expressed transcript | 41.5 | 42.75 | 40.66 | 41.9 |
| 947773 | 23S ribosomal RNA of rrnD operon | 53.4 | 51.29 | 53.05 | 55.57 |
| 50652 | prostate cancer associated | 40.88 | 41.21 | 41.12 | 40.3 |
| 5820 | Pvt1 oncogene |  | 46.8 | 47.85 | 48.95 |
| 55000 | taurine up-regulated 1 | 42.05 | 40.51 | 43.37 | 42.26 |
| 5802729 | prl45 | 30.26 | 29.58 | 32.2 | 29.21 |
| 66961 | nuclear paraspeckle assembly transcript 1 | 47.26 | 47.66 | 46.68 | 47.75 |
| 10037957 |  |  |  |  |  |
| 1 | BACE1 antisense RNA | 48 | 49.63 | 44.16 | 49.81 |
| 10115464 | long intergenic non-protein coding RNA,muscle |  |  |  |  |
| 4 | differentiation | 40.92 | 40.69 | 41.72 | 40.34 |
| 55539 | KCNQ1 downstream neighbor | 60.97 | 60.86 | 59.56 | 62.37 |
| 26534084 | Dpse\GA32664 | 35.23 | 35.4 | 35.27 | 34.99 |
| 10537381 |  |  |  |  |  |
| 8 | long intergenic non-protein coding RNA 1827 | 40.44 | 40.86 | 39.87 | 40.59 |
| IV. | Table: The overall and position specific GC content of noncoding regions |  |  |  |  |

## REFERENCES

1. Larry Snyder, and Wendy Champness. Molecular Genetics Of Bacteria. 3rd ed. 2003. Print.

2. Hershberg, Ruth, and Dmitri A. Petrov. “Selection On Codon Bias”. The Annual Review of Genetics 42 (2008): 288. Web. 13 June 2017.

3. Crick, F. H. C. et al. “General Nature Of The Genetic Code For Proteins”. Nature 192.4809 (1961): 1227–1232. Web.

4. Sharp, Paul M., and Wen-Hsiung Li. “The Codon Adaptation Index-A Measure Of Directional Synonymous Codon Usage Bias, And Its Potential Applications”. Nucleic Acids Research 15.3 (1987): 1281–1295. Web.

5. Sharp, Paul M. et al. “Codon Usage Patterns In Escherichia Coli, Bacillus Subtilis, Saccharomyces Cerevisiae, Schizosaccharomyces Pombe, Drosophila Melanogaster and homo Sapiens; A Review Of The Considerable Within-Species Diversity”. Nucleic Acids Research 16.17 (1988): 8207–8211. Web.

6. Grantham, R. et al. “Codon Catalog Usage And The Genome Hypothesis”. Nucleic Acids Research 8.1 (1980): 197–197. Web.

7. Grantham, R. et al. “Codon Catalog Usage Is A Genome Strategy Modulated For Gene Expressivity”. Nucleic Acids Research 9.1 (1981): 213–213. Web.

8. Ikemura, Toshimichi. “Correlation Between The Abundance Of Escherichia Coli Transfer Rnas And The Occurrence Of The Respective Codons In Its Protein Genes: A Proposal For A Synonymous Codon Choice That Is Optimal For The E. Coli Translational System”. Journal of Molecular Biology 151.3 (1981): 389–409. Web.

9. Ikemura, Toshimichi. “Correlation Between The Abundance Of Yeast Transfer Rnas And The Occurrence Of The Respective Codons In Protein Genes”. Journal of Molecular Biology 158.4 (1982): 573–597. Web.

10. Sørensen, Michael A., C.G. Kurland, and Steen Pedersen. “Codon Usage Determines Translation Rate In Escherichia Coli”. Journal of Molecular Biology 207.2 (1989): 365–377. Web.

11. Parker, J., T. C. Johnston, P. T. Borgia, G. Holtz, E. Remaut et al., “Codon Usage and Mistranslation - In Vivo Basal Level Misreading Of The MS2 Coat Protein Message”. Journal of Biological Chemistry 258 (1983): 7–12. Print.

12. Bennetzen, J. L. & Hall, B. D “Codon Selection In Yeast”. Journal of Biological Chemistry 257 (1982): 3026–3021. Print.

13. Sharp, P. M., L. R. Emery, and K. Zeng. “Forces That Influence The Evolution Of Codon Bias”. Philosophical Transactions of the Royal Society B: Biological Sciences 365.1544 (2010): 1203–1212. Web.

14. Kimura, Motoo. “A Simple Method For Estimating Evolutionary Rates Of Base Substitutions Through Comparative Studies Of Nucleotide Sequences”. Journal of Molecular Evolution 16.2 (1980): 111–120. Web.

15. Plotkin, Joshua B., and Grzegorz Kudla. “Synonymous But Not The Same: The Causes And Consequences Of Codon Bias”. Nature Reviews Genetics 12.1 (2010): 32–42. Web.

16. Comeron, J. M., Kreitman, M. & Aguade, M. “Natural Selection On Synonymous Sites Is Correlated With Gene Length And Recombination In Drosophila.”. Genetics 151 (1999): 239–249. Print.

17. Marais, G., D. Mouchiroud, and L. Duret. “Does Recombination Improve Selection On Codon Usage Lessons From Nematode And Fly Complete Genomes”. Proceedings of the National Academy of Sciences 98.10 (2001): 5688–5692. Web.

18. Galtier, Nicolas. “Gene Conversion Drives GC Content Evolution In Mammalian Histones”. Trends in Genetics 19.2 (2003): 65–68. Web.

19. Marais, G., D. Mouchiroud, and L. Duret. “Neutral effect of Recombination in base composition of Drosophila”. Genetics Research 81(2003): 79–87. Web.

20. Behura, Susanta K., and David W. Severson. “Codon Usage Bias: Causative Factors, Quantification Methods And Genome-Wide Patterns: With Emphasis On Insect Genomes”. Biological Reviews 88.1 (2012): 49–61. Web.

21. Shannon, C. E. “A Mathematical Theory Of Communication”. Bell System Technical Journal 27.4 (1948): 623–656. Web.

22. Adami, Christoph. “Information Theory In Molecular Biology”. Physics of Life Reviews 1.1 (2004): 3–22. Web.

23. Zeeberg, B. “Shannon Information Theoretic Computation Of Synonymous Codon Usage Biases In Coding Regions Of Human And Mouse Genomes”. Genome Research 12.6 (2002): 944–955. Web.

24. “Codon Usage”. Bioinformatics.org. N.p., 2017. Web. 14 June 2017.

25. “Random DNA Generator”. Faculty.ucr.edu. N.p., 2017. Web. 14 June 2017.

26. Wan, Xiu-Feng et al. “Quantitative Relationship Between Synonymous Codon Usage Bias And GC Composition Across Unicellular Genomes”. BMC Evolutionary Biology (2004): 4–19. Print.

27. Galtier, N et al. “GC-Content Evolution In Mammalian Genomes: The Biased Gene Conversion Hypothesis”. Genetics Society of America 159 (2001): 907. Print.

28. Gerton, J. L. et al. “Global Mapping Of Meiotic Recombination Hotspots And Coldspots In The Yeast Saccharomyces Cerevisiae”. Proceedings of the National Academy of Sciences 97.21 (2000): 11383–11390. Web.

29. Brown, Thomas C., and Josef Jiricny. “A Specific Mismatch Repair Event Protects Mammalian Cells From Loss Of 5-Methylcytosine”. Cell 50.6 (1987): 945–950. Web.

30. C A, Bill, and Duran W A. “Efficient Repair Of All Types Of Single-Base Mismatches In Recombination Intermediates In Chinese Hamster Ovary Cells. Competition Between Long-Patch And G-T Glycosylase-Mediated Repair Of G-T Mismatches”. Genetics 149 (1998): n. pag. Print.

31. Kimura, MOTOO. “Evolutionary Rate At The Molecular Level”. Nature 217.5129 (1968): 624–626. Web.

32. Crick, F.H.C. “Codon—Anticodon Pairing: The Wobble Hypothesis”. Journal of Molecular Biology 19.2 (1966): 548–555. Web.

